# The Gut Microbiome and Butyrate Differentiate *Clostridioides difficile* Colonization and Infection in Children

**DOI:** 10.1101/2025.04.16.648821

**Authors:** Maribeth R. Nicholson, Siyuan Ma, Britton A. Strickland, Mia Cecala, Lisa Zhang, Seth Reasoner, Emma R. Guiberson, Matthew J Munneke, Meghan H. Shilts, Eric P. Skaar, Suman R. Das

**Affiliations:** Division of Pediatric Gastroenterology, Hepatology, and Nutrition, Department of Pediatrics, Vanderbilt University Medical Center, Nashville, TN, USA; Department of Pathology, Microbiology, and Immunology, Vanderbilt University Medical Center, Nashville, TN, USA; Division of Molecular Pathogenesis, Department of Pathology, Microbiology & Immunology, Vanderbilt University Medical Center, Nashville, TN, USA; Department of Chemistry and Biochemistry, Middlebury College, Middlebury, VT; Division of Infectious Diseases, Vanderbilt University Medical Center, Nashville, TN

**Author notes:** Corresponding authors: 1. Suman R. Das PhD, Address: Division of Infectious Diseases, Vanderbilt University Medical Center, 1211 21st Avenue South, S2108 Medical Center North, Nashville, TN 37232, 2. Maribeth Nicholson, MD MPH, Maribeth, Address: DOT 10^th^ floor, 2200 Children’s Avenue, Nashville, TN 37232. Denotes co-first authors as contributed equally to the work (Siyuan Ma, Britton A. Strickland).

**Keywords:** toxins, pediatric, carriage, metabolite

## Abstract

**Background and Aims:** Symptomatic *Clostridioides difficile* infection (CDI) can cause significant morbidity and mortality. Conversely, patients can be colonized with toxigenic *C. difficile* in the absence of symptoms, termed asymptomatic colonization. We previously demonstrated that the presence and function of *C. difficile* toxins do not differentiate between asymptomatic colonization and CDI in children, suggesting the influence of other factors. This study aimed to interrogate the intestinal microbiome and butyrate in stool samples from children with CDI and asymptomatic colonization.

**Methods:** Design: Case-control study

Setting: Tertiary care children’s hospital

Participants and measures: Asymptomatic children had stool tested for *C. difficile* by nucleic-acid amplification-based testing (NAAT) and were considered colonized if positive (N=50). Residual stool was also obtained from symptomatic children who tested positive for *C. difficile* by NAAT (N=55). The microbiome was assessed via 16S rRNA sequencing and butyrate via liquid chromatography-mass spectrometry.

**Results:** Compared to clinical co-variates and comorbidities, *C. difficile* symptom status (i.e., asymptomatic colonization versus symptomatic CDI) demonstrated the strongest differential abundance association on gut microbes. Symptomatic CDI was associated with increased abundance of *Escherichia/Shigella* (Benjamini-Hochberg adjusted q=3.94×10^−5^), *Haemophilus* (q=0.022), and *Gemella* (q=0.085), and depleted abundance of gut commensals such as *Faecalibacterium* (q=0.041), *Blautia* (q=0.041), and *Bifidobacterium* (q=0.063). We also observed depletion in the abundance of microbial butyrate producers and fecal butyrate in participants with symptomatic CDI versus asymptomatic colonization.

**Conclusion:** The gut microbiota and butyrate differ between participants with asymptomatic *C. difficile* colonization and symptomatic CDI, suggesting their potential role in symptom development.

## INTRODUCTION

*Clostridioides difficile* is a gastrointestinal pathogen with a global health impact and is the leading cause of antibiotic-associated diarrhea.^1^ It has demonstrated a rising incidence in both children and adults and can be associated with substantial morbidity and mortality.^2^ Alternatively, due to unclear mechanisms, *C. difficile* can colonize the intestinal tract without causing symptomatic disease, referred to as asymptomatic colonization. Children with the highest rates of asymptomatic *C. difficile* colonization include those with cancer, inflammatory bowel disease (IBD), and cystic fibrosis (CF), who are also at risk for diarrhea from a variety of alternative causes.^3–5^ No testing algorithm has reliably differentiated symptomatic *C. difficile* infection (CDI) from those with asymptomatic colonization with diarrhea from an alternative source. Although classically used in diagnostic testing algorithms, our previous work demonstrated that the presence and activity of *C. difficile*-associated toxins are unable to differentiate symptomatic CDI from asymptomatic colonization in a pediatric cohort.^6^ In addition, the same *C. difficile* strains and bacterial burden associated with asymptomatic colonization in some patients can be associated with severe disease in others.^7^ This, therefore, highlights an important question; if the presence of *C. difficile* and toxins are similar between those with asymptomatic colonization and symptomatic disease, what then differs? This question is important for two reasons: 1) to improve diagnostic testing and its interpretation and 2) to understand protective mechanisms that influence the pathophysiology of *C. difficile*.

Despite pathogenesis tightly linked to the intestinal microbiome,^8^ little data have been published on alterations in the intestinal microbiome and associated metabolites in pediatric patients with CDI versus asymptomatic colonization. Previously published studies have been limited by small sample sizes^9,10^ or performed exclusively in adults.^11,12^ A single adult study of eight patients demonstrated decreased alpha and beta diversity in patients with symptomatic CDI compared to colonized patients.^11^ Alternatively, Crobach et al. found no differences in alpha and beta diversity in adult patients with CDI and asymptomatic colonization. Still, they identified that patients with symptomatic CDI had a significantly higher relative abundance of *Bacteroides* and *Veillonella* than colonized patients.^12^

Microbially mediated metabolites may also play a critical role in preventing symptomatic CDI. Butyrate, a short-chain fatty acid produced through the metabolism of dietary fibers, has demonstrated the potential to reduce intestinal inflammation and mitigate the development of *C. difficile*-induced colitis in a murine model.^13^

Here, using a well-characterized cohort of pediatric patients, we compared the intestinal microbiome and butyrate levels in children with symptomatic CDI and asymptomatic colonization to better understand the pathophysiology of this important bacterium.

## MATERIALS AND METHODS

### Study Design

Pediatric participants, ages 12 months through 18 years, were prospectively enrolled from July 2017 through December 2019 at Monroe Carell Jr. Children’s Hospital at Vanderbilt after informed parental consent and patient assent when applicable. The Vanderbilt Institutional Review Board approved the study. Thorough medical histories were obtained on all participants, including past hospitalizations, surgeries, and medications received 30 days before enrollment and confirmed by medical record review. Data were kept confidential using a REDCap database (REDCap software, Vanderbilt University).^14^

### Asymptomatic Colonized Cohort

Asymptomatic children between 12 months and 18 years of age with a diagnosis of cancer, cystic fibrosis (CF), or inflammatory bowel disease (IBD) were eligible for enrollment. To be considered asymptomatic, participants had to be without diarrhea (Bristol stool type 6 or 7)^15^ and not undergoing active testing or treatment for *C. difficile.* Patients were recruited during outpatient visits or hospitalizations, and a stool sample was obtained. At the time of processing, an aliquot of stool underwent testing for *C. difficile* by nucleic-acid amplification-based testing (NAAT) in the clinical laboratory. If positive by NAAT, the child was considered colonized.

### Symptomatic Cohort

Symptomatic children with diarrhea (≥three unformed stools in 24 hours and an acute change in symptoms) between 12 months and 18 years of age who underwent clinical laboratory testing and tested positive for *C. difficile* by NAAT were enrolled. This included previously healthy children and those with additional comorbidities. After consent, residual stool from the clinical laboratory was collected and stored in the research laboratory. Enrollment schemes are compared in **Table 1**.

**Table 1.** Comparison of enrollment scheme.

|  | Asymptomatic Cohort | Symptomatic Cohort |
| --- | --- | --- |
| Cohort Definition | • Diagnosis of IBD, CF, or Cancer<br>• Formed stool<br>• Not being tested or treated for CDI<br>• Positive *C. difficile* NAAT test | • All comorbidities or healthy<br>• Positive *C. difficile* NAAT test<br>• $\geq 3$ unformed BM per 24 hours<br>• Acute change in symptoms<br>• Residual stool collected from VUMC laboratory |
| Cohort Participants | N = 50 | N = 55 |
| Sample Analysis | ELISA for toxin presence<br>Vero cell rounding for toxin function<br>16S rRNA sequencing for microbiome assessment<br>LC-MS for fecal butyrate quantification |  |

### Sample Processing and Clinical Testing

Stools were refrigerated immediately after collection and stored at −80°C after aliquoting. NAAT was performed in the hospital clinical laboratory for both asymptomatic and symptomatic participants using Illumigene *C. difficile* assay (ARUP laboratories), a polymerase chain reaction test to detect the *C. difficile* gene *tcdB* encoding Toxin B (Sensitivity 90%, Specificity 96%).^16,17^ Toxin presence was determined in the research laboratory in duplicate with enzyme-linked immunoassay testing using Premier® Toxins A and B from Meridian Biosciences per manufacturer recommendations (Sensitivity 80.8%, Specificity 97.5%).^16,18^ Toxin function was determined by Vero cell rounding, which was performed as previously described.^19^

### DNA Extraction and Microbiome Sequencing

Genomic DNA extraction (Qiagen DNeasy PowerSoil Kit), targeted PCR (V4 region of 16S rRNA gene^20^), and paired-end sequencing (Illumina MiSeq 2×250bp) was performed at Vanderbilt University Medical Center as previously described^21^.

### 16S rRNA Data Processing

16S rRNA sequences grouped into amplicon sequence variants (ASVs) and assigned taxonomy using the dada2 pipeline (available at: https://benjjneb.github.io/dada2/tutorial.html, last accessed September, 2020)^22^ and the SILVA reference database.^23^ Suspected contaminants in the negative controls were removed using the R package decontam^24^. The R package phyloseq^25^ was also used to facilitate data processing.

### Statistical Analysis

Most statistical analyses were conducted under the open source R computing environment. Alpha- and beta-diversity metrics were calculated with the R package *vegan*^26^ at the ASV level. Difference associations in alpha-diversity metrics are based on non-parametric Kruskal-Wallis tests. Significant association between beta-diversity (Bray-Curtis) was assessed using PERMANOVA with 99,999 permutations. Significant associations between metadata and bacterial taxa were evaluated at the phylum, genus, and ASV levels. These were based on microbiome multivariable linear regression as implemented in the R package MaAsLin2.^27^ We primarily adjusted for covariates including demographics (age, gender), comorbidities, and exposure history (antibiotics, acid blockers, hospitalization, surgery), but conducted additional sensitivity analyses where (1) we stratified by comorbidity groups to ensure that the associations were consistent, and (2) we adjusted for different subsets of covariates, including both unadjusted analyses and analyses adjusting for only age, gender, and comorbidity, again to ensure that findings were consistent. Multiple testing across different microbial taxa were corrected using the Benjamini-Hochberg procedure. Findings were further filtered to remove microbes that were present in five or fewer samples.

Associations were considered significant if the p- or q-value (as appropriate) was <0.05 and <0.10 respectively.

### Butyrate measurements by liquid chromatography-mass spectrometry

#### Butyrate extraction and derivatization

Human fecal matter was aliquoted, weighed, diluted in MeOH/H2O (1:5), and homogenized using a bead beater. Samples were centrifuged to remove remaining debris, and resulting supernatants were transferred to Eppendorf tubes and stored at −20 °C until analysis. Butyrate was derivatized using dansylhydrazine and 1-Ethyl-3-(3-dimethylaminopropyl)carbodiimide (EDC) as a carboxyl activating agent. 10 µL of fecal extracts were spiked with isotopically-labeled butyrate (butyrate-d_5_) and derivatized in a 50 mM sodium phosphate buffer (pH = 4.0) in H_2_O/DMSO (2:1) with 12.5 mg/mL dansylhydrazine and 12.5 mg/mL EDC. Samples were incubated at room temperature for 2 hours, then 100 mM NaOH was added to solution. Dansylated derivatives were then extracted using 750 µL of ethyl acetate, and the organic layer was collected and transferred to a new tube, dried down using nitrogen gas, and reconstituted in 150 µL of 50:50 acetonitrile:water. Calibration standards were prepared in water and derivatized and extracted using the same procedure. Due to the possible influence of water content, a subset of samples was lyophilized, and extraction and derivatization were performed to ensure that trends were maintained.

#### Liquid chromatography-tandem mass spectrometry analysis

Butyrate measurements and analysis were performed using a Thermo TSQ Quantum mass spectrometer operated in positive ionization mode coupled to a Thermo Surveyor HPLC pump and autosampler. For liquid chromatography, a reverse phase method was utilized using a Acquity BEH C18 analytical column. Mobile phases were made up with 0.2% acetic acid, 15 mM ammonium acetate in either H_2_O/ CH_3_CN 9:1 (A) or CH_3_CN/ CH_3_OH/ H_2_O 90:5:5 (B). The following gradient conditions were used: 0-1 min 0% B, 1-8 min 0-100% B, 8-10 min 100% B, 10-10.5 min 100-0% B, 10.5-15 min 0% B with a flow rate of 300 µL/min for a total gradient time of 15 minutes. Eluent between 0 and 2 minutes were diverted to waste. Samples were maintained at 5 °C in the autosampler, and the column was held at 50 °C. Injection volume was 10 µL. Quantitation was based on single reaction monitoring detection of butyrate dansylated analogues: butyrate, *m/z* 336 171, CE 30; butyrate -d_5_ *m/z* 341 170, CE 30. Data analysis was done using Thermo-Finnigan Xcaliber. Calibration curves were constructed by plotting peak area ratios against analyte concentrations, for a series ranging from 0.01 to 100 nmol butyrate and used to determine pmol concentrations. Samples were then normalized by weight for final pmol/mg values. Statistical analyses were performed using Mann-Whitney U-tests through GraphPad Prism.

## RESULTS

### Study population

The study population included 50 pediatric participants with asymptomatic *C. difficile* colonization and 55 children with symptomatic CDI (**Table 1**). The median age of the cohort was ten years (interquartile range (IQR) 5-15), and 55% of the cohort were male. The asymptomatic colonized cohort included 32 (64%) children with CF, 8 (16%) children with IBD, and 10 (20%) children with cancer. Patients with asymptomatic colonization and symptomatic CDI were compared (**Table 2**). Symptomatic children were older (median 13 [IQR 5-16] vs 9 [4-12] years, *P*=0.025) and had a different comorbidity profile based on enrollment strategy (*P*<0.001). Asymptomatic colonized children were more likely to report acid blocker use in the 30 days before enrollment (76% vs 42%, *P*<0.001). Antibiotic use between cohorts was similar (54% vs. 67%, *P*=0.23). C-reactive protein (CRP) values were higher in the symptomatic cohort (median 9.5 [IQR 3.4-35.8] vs 2.6 [IQR 0.2-7.4], P=0.047), but the remainder of laboratory values were not significantly different.

**Table 2.**
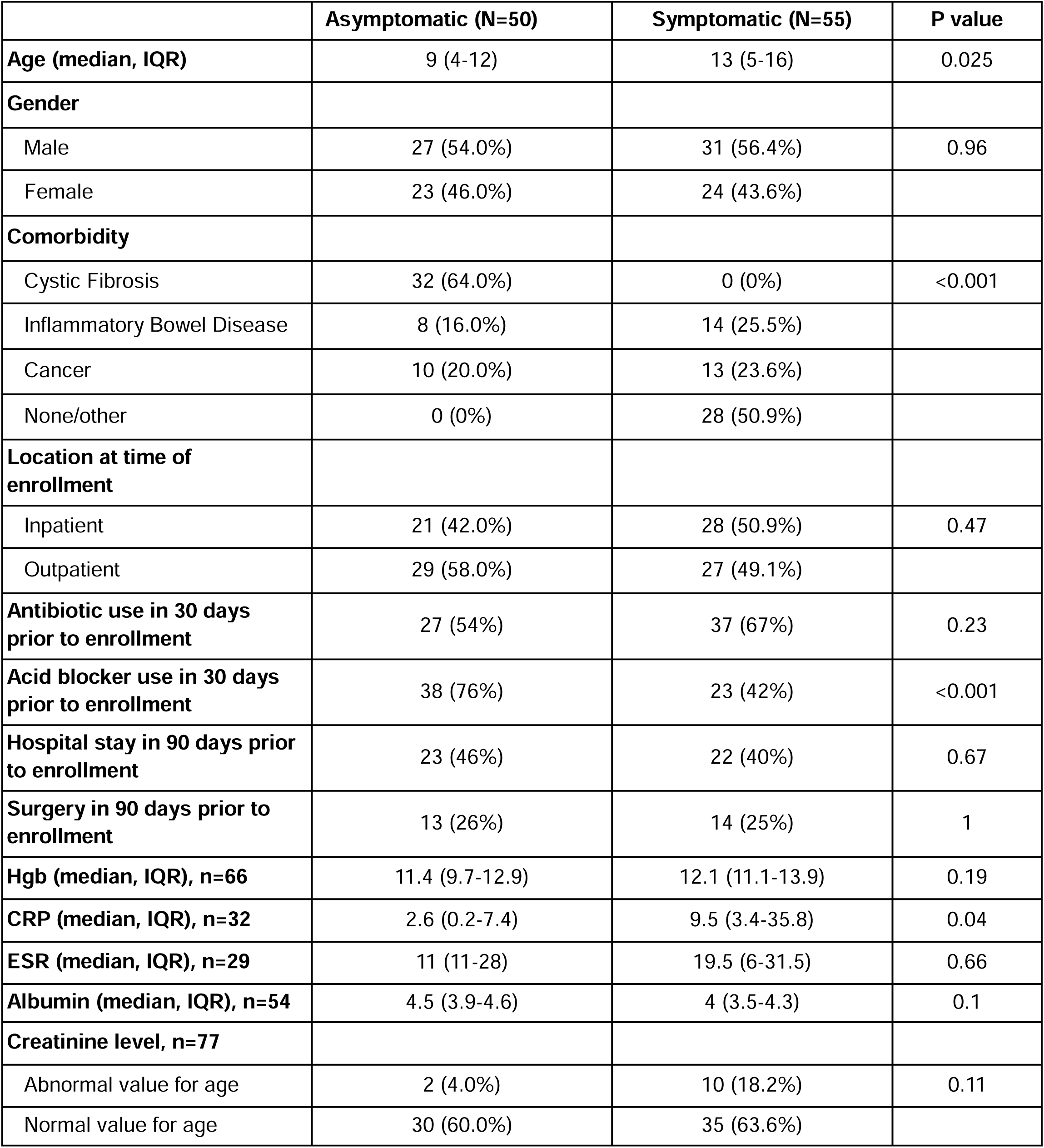
Demographic and clinical variables of patient cohorts. Continuous variables are expressed as median (interquartile range [IQR]). Categorical and dichotomous variables are expressed as n and proportion of the respective group’s total. Statistical analyses were conducted by logistic regression models in R statistical software. Abbreviations: Hgb: hemoglobin; CRP: C-reactive protein; ESR: erythrocyte sedimentation rate.

### ELISA and Vero Cell Rounding results do not correspond to symptom status

ELISA and Vero Cell rounding were used to assess *C. difficile* toxin presence and function in samples that tested positive for *C. difficile* via NAAT. A portion of this cohort was previously published.^19^ In symptomatic cases, 60.7% of the samples were positive for Toxin A/B ELISA versus 39.3% of those with asymptomatic colonization (P=0.32) **(Figure S1A).** In addition, 50% of symptomatic patients tested positive for Vero cell rounding toxicity assessment versus 50% of those with asymptomatic colonization (P=0.88) **(Figure S1A).** These results indicate that toxin-based tests cannot differentiate between symptomatic and asymptomatic colonized pediatric patients (hereby referred to as *C. difficile* symptom status). Both positive ELISA and Vero cell rounding were associated with an increased abundance of *C. difficile* (Figure S1B, Figure S1C) regardless of symptom status. Analysis of richness and alpha diversity did not find any significant differences between samples testing positive and negative for ELISA and Vero cell rounding (**Figure S1D**).

### Comparison of gut microbiome between asymptomatic C. difficile colonized and symptomatic patients

Microbiome profiling was performed using 16S rRNA amplicon sequencing of the V4 gene region. For diversity and differential taxa testing, a sample read cutoff of >1000/sample was implemented with the lowest library size of 1533 reads and 90.7% samples >10,000 reads. We then examined the association of diversity and taxa between symptomatic and asymptomatic colonized patients, as well as a subset of patients with IBD and cancer.

We observed substantial differences in overall microbiome community diversity when comparing cohorts by *C. difficile* symptom status (**Figure 1**, **Figure S2**). Asymptomatic colonized patients had a higher alpha diversity/evenness based on Shannon (Mann-Whitney U p=0.02352) and Simpson (Mann-Whitney U p=0.00515) indices (**Figure 1A)**. Beta diversity was performed using Bray-Curtis dissimilarity at the ASV level and showed a significant separation of community composition between asymptomatic colonized and symptomatic patients (**Figure 1B**; PERMANOVA p<10^−5^). Such differences in microbiome composition are directly visualized with a stacked barplot of dominant gut microbiome genera in **Figure S2**. Additionally, beta diversity analysis with respect to covariates in this study revealed the impact of various exposures (antibiotic use, acid blocker use, hospital stay, and surgery) as well as comorbidities on the intestinal microbiome, suggesting the importance of adjusting for their potential confounding effects (**Figure S3**).

**Figure 1.**
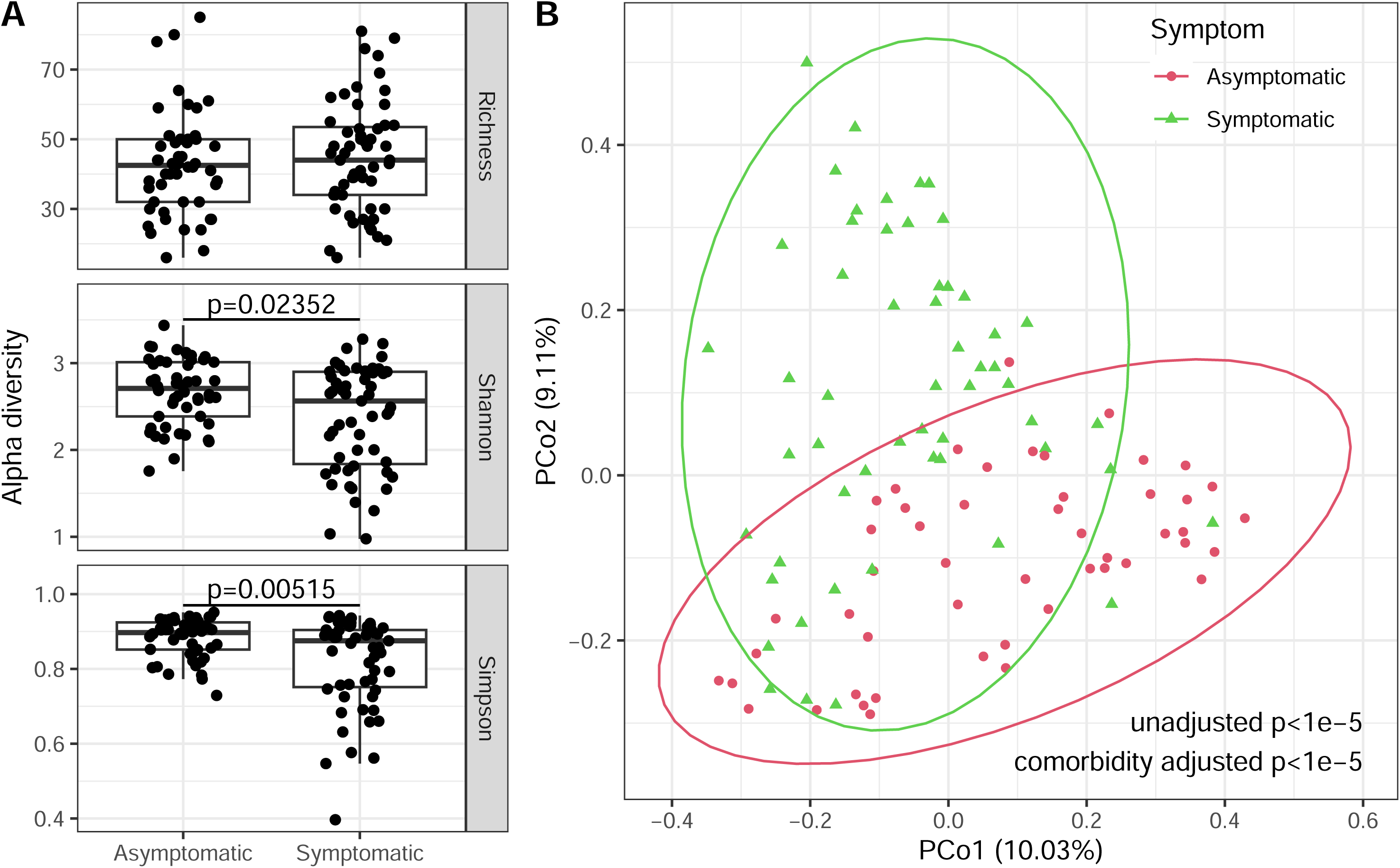
Gut microbiome diversity is significantly associated with symptomatic vs. asymptomatic difference. **A.** Alpha diversity indices, including Shannon diversity and Simpson index, are significantly decreased in symptomatic samples, suggesting that symptomatic subjects have less diverse gut microbiome compared to asymptomatic. p-values are based on nonparametric Kruskal-Wallis tests. **B.** Beta diversity analysis (Bray-Curtis dissimilarity) indicates significant difference between the gut microbiome profile of symptomatic compared to asymptomatic subjects. p-values are based on PERMANOVA analyses with 99999 permutations.

Individual microbial taxa were strongly associated with *C. difficile* symptom status. The gut microbiome of patients with symptomatic CDI was marked by an expansion of dysbiotic microbes and a depletion of commensals (**Figure 2A, Figure S4**).

**Figure 2.**
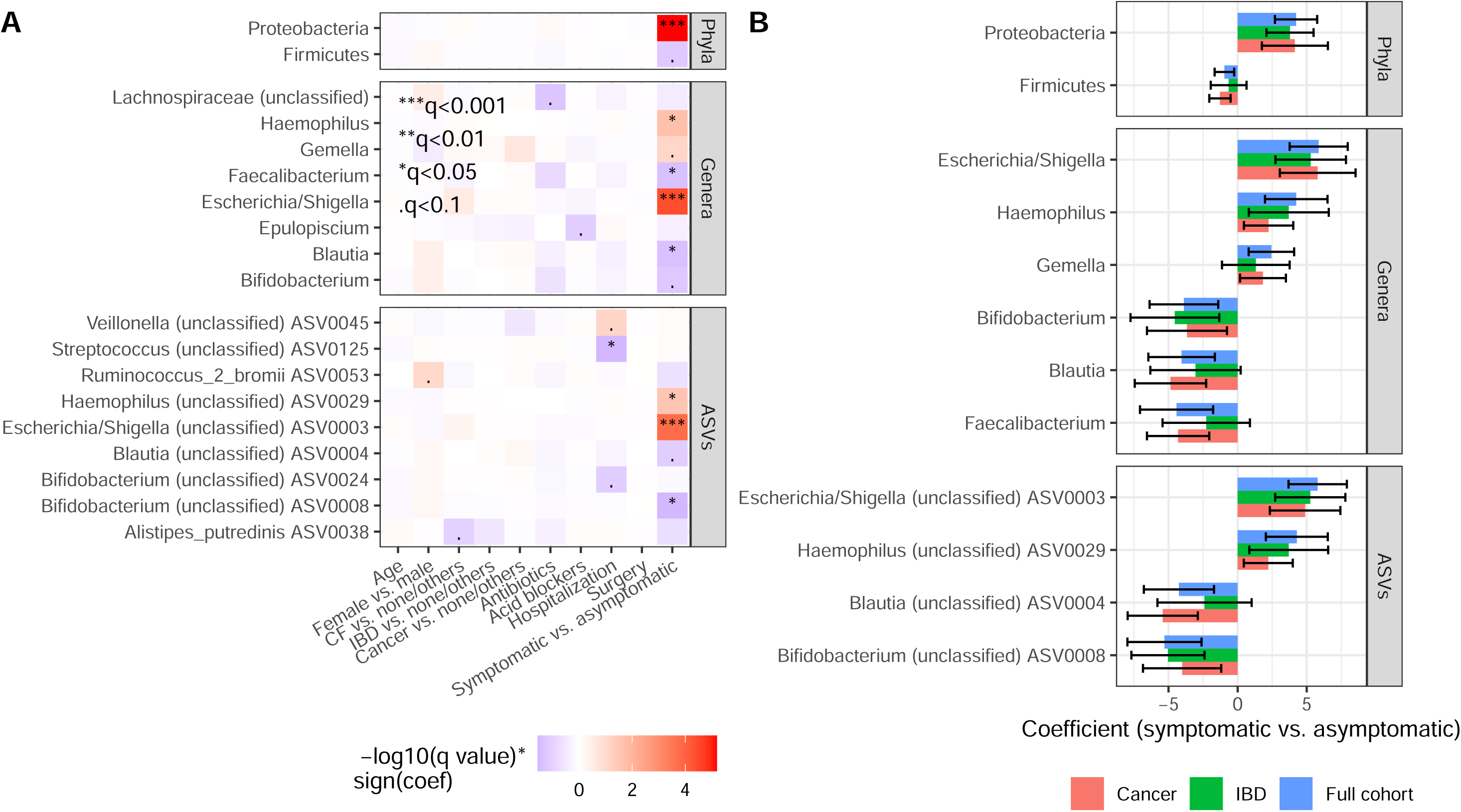
Abundance of gut microbes are significantly associated with symptomatic versus asymptomatic infections; such difference is consistent across comorbidity groups. **A.** Microbes with significant differential abundance in association with the outcome (infection symptom) and covariate variables in this study, including demographics (age, gender), comorbidity, and prior intervention history (antibiotics, acid blocker, hospitalization, and surgery). We report associated microbial taxa at the phylum, genus, and ASV level. Differential abundance analyses are performed with multiple linear regression modeling (MaAsLin2) and adjusted for multiple hypothesis testing burden with the Benjamini-Hochberg method. **B.** Identified microbes associated with infection symptoms (q < 0.1 from panel A) also have consistent differential abundant effects when analysis is restricted to comorbidity subgroups (cancer, inflammatory bowel disease). x-axis represents differential abundant regression coefficient (log fold change). Error bars indicate 95% confidence intervals.

Compared to all the adjusted covariates, *C. difficile* symptom status demonstrated the strongest and most significant differential abundance association on gut microbes (significance determined with Benjamini-Hochberg adjusted q < 0.1), with six significantly associated microbes at the genus level (**Figure 2A, Figure S4, Table S1**). Symptomatic CDI was associated with increased abundance of *Escherichia/Shigella* (Benjamini-Hochberg adjusted q=3.94×10^−5^), *Haemophilus* (q=0.022),), and *Gemella* (q=0.085) and depleted abundance of common gut commensals such as such as *Faecalibacterium* (q=0.041), *Blautia* (q=0.041), and *Bifidobacterium* (q=0.063). These associations were confirmed at the ASV level (representing species or sub-species level resolution), whereby members of the identified genera (*Hemophilus*, *Escherichia/Shigella*, *Blautia*, and *Bifidobacterium*) demonstrated similar associations. At the phylum level, patients with symptomatic CDI had a significant expansion in Proteobacteria gut microbes and depletion of Firmicutes. Some gut microbes were also significantly associated with exposure history, including depletion of an unclassified member of *Lachnospiraceae* with antibiotic use (q=0.057) and depletion of *Epulopiscium* with prior hospital stay (q=0.081). Our findings indicate that the gut microbiome significantly differs between asymptomatic colonized patients versus those with symptomatic CDI, after adjusting for demographics, exposure history, and comorbidities.

To ensure that the observed associations were not due to confounding by comorbidity effects, we conducted a detailed secondary analysis to examine the differential abundant effects of the microbes associated with symptom status in the entire cohort. This was performed separately in participants with IBD and cancer, which were represented in both symptomatic CDI and asymptomatic colonization cohorts (**Figure 2, Table S2**). Again, we performed multilinear regression by adjusting for demographic and intervention history variables. The microbes had consistent differential abundant effects in both patient cohorts, solidifying that our observed associations were indeed driven by symptom status and not comorbidity profile.

Lastly, sensitivity analysis revealed that the association between gut microbes and infection symptom status were consistent when different sets of covariates were adjusted in the analysis (**Figure S5, Table S3**). Our findings support that the association of gut microbes with symptom status were not sensitive to patient comorbidities or antibiotic exposure history.

### Butyrate and butyrate-producing microbes are potential mitigators of CDI

It has been reported that butyrate and butyrate-producing gut microbes are significantly depleted in CDI patients,^13,28^ and indeed may function as therapeutics to mitigate CDI. However, previous studies have not differentiated between symptom phenotypes of *C. difficile.* We, therefore, investigated differences in butyrate and butyrate-producing organisms in children with symptomatic CDI and asymptomatic colonization.

We observed strong depletion in both the abundance of microbial butyrate producers and butyrate concentration in symptomatic patients of our cohort (**Figure 3**). Given major butyrate producers as compiled by Singh et al.,^29^ we observe that their total abundance was significantly depleted in symptomatic patients, adjusting again for demographics, exposure history, and comorbidities (**Figure 3A**). Further investigation of individual butyrate-producing microbes at both the genus and ASV level revealed that most demonstrated a negative association with symptomatic CDI (**Figure 3B**), although most did not reach statistical significance except for *Blautia* (q=0.041) and *Faecalibacterium* (q=0.041; **Table S4**), potentially limited by the study sample size. Additionally, the decreased abundance of butyrate-producing microbes in CDI was mirrored by depletion of butyrate stool concentration in the symptomatic CDI subjects based on LC-MS (**Figure 3C**). Due to the potential differences in water content among samples, a subset underwent lyophilization, and LC-MS was repeated. The trend was maintained, with asymptomatic colonized participants harboring ∼10-fold higher butyrate concentrations than participants with symptomatic CDI (**Figure S6**). Overall, our results strongly support previous findings on the protective effect of butyrate and butyrate-producing bacteria and suggest the role of butyrate in the prevention of symptoms in those with asymptomatic colonization.

**Figure 3.**
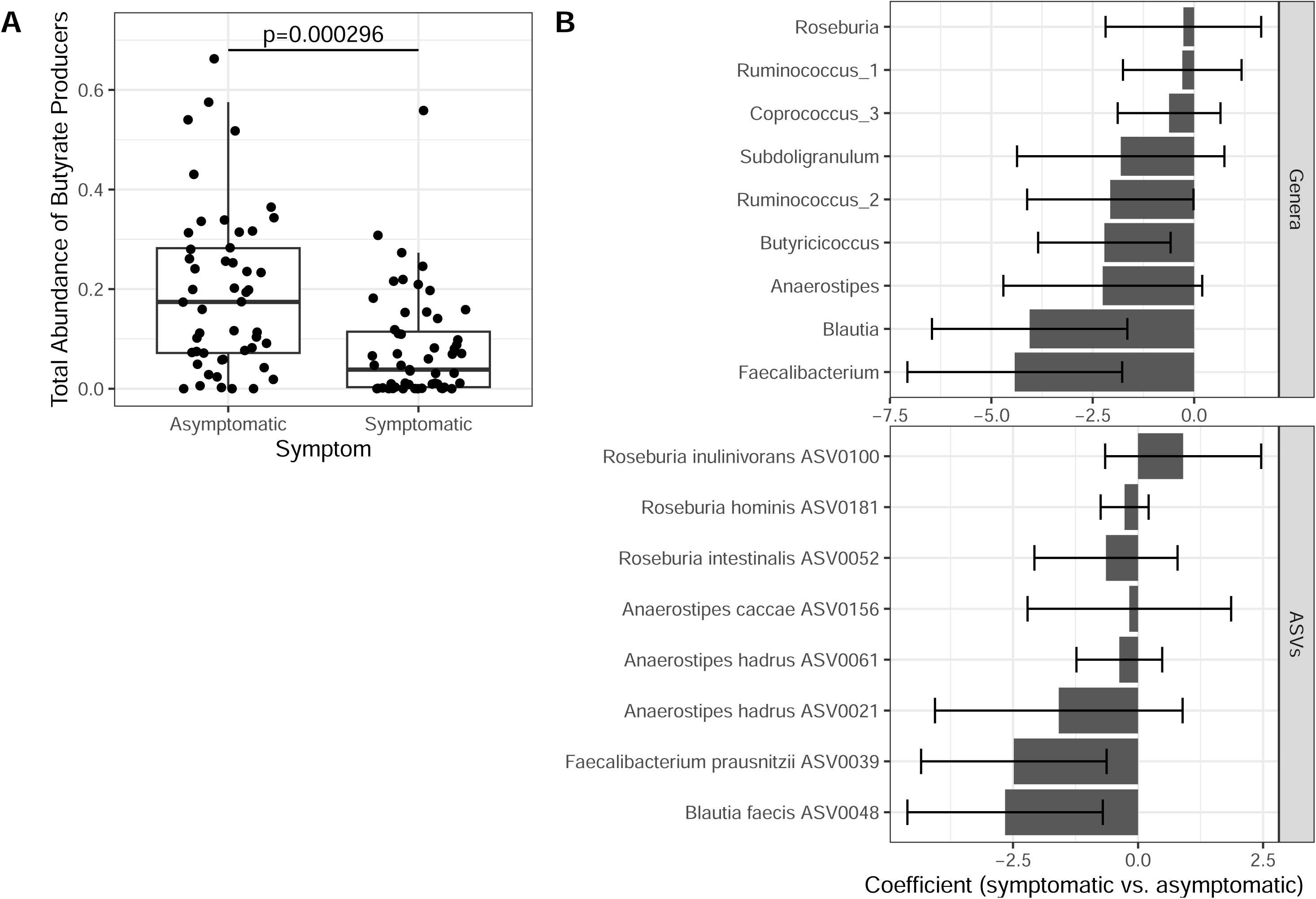
Butyrate and butyrate producing bacteria are likely mitigators of C. difficile infection toxicity. **A.** Total abundance of major butyrate producing gut genera (as summarized in {36713166}) is significantly decreased in symptomatic compared to asymptomatic patients. This is congruent with per-taxon findings, where most butyrate producing taxa (genera and ASVs, the latter representing species resolution) had reduced abundance in symptomatic patients (**B**). For **A** and **B** statistical analysis is performed with same multiple linear modeling as in Figure 3A and adjusted for the same set of covariates. For **B** x-axis represents differential abundant regression coefficient (log fold change). Error bars indicate 95% confidence intervals. **C.** Patients asymptomatically colonized by *C. difficile* show higher levels of fecal butyrate than symptomatic patients. Quantification of butyrate by LC-MS/MS in human fecal matter from patients asymptomatically or symptomatically colonized with *C. difficile*. N = 40 per group, outliers removed using ROUT test with Q < 0.2%. Data are represented as mean±standard error to the mean. Statistical significance was assessed using a Mann-Whitney U-test. ****p<0.0001.

## DISCUSSION

Infection with *Clostridioides difficile* is a threat to public health and associated with significant morbidity and mortality.^1^ Although a toxin-mediated disease, there are reports of patients with a high abundance of both *C. difficile* and its associated toxins who remain without clinical symptoms, deemed asymptomatic colonization.^30^ It is well established that these patients may serve as an infectious vector and create diagnostic challenges if they develop diarrhea from an alternative cause.^3,7^ However, they also may provide significant insights into factors that protect against the development of symptomatic CDI and warrant targeted study.

Our prior work has attempted to determine the role of toxin presence and activity in pediatric patients with asymptomatic colonization compared to those with CDI. Toxins are a necessary component of symptomatic CDI through their ability to cause inflammation and cell damage to the colonic epithelium, resulting in fluid secretion and diarrhea. Contrary to the association of toxins with inflammation, our prior cohort of asymptomatic colonized pediatric patients demonstrated similar toxin presence and activity compared to those with symptomatic CDI.^31^ Notably, many children with active toxins, based on Vero-cell rounding, remained well. Similarly, in this study that expands upon the prior cohort, the presence and activity of toxins could not differentiate between those with asymptomatic colonization and symptomatic CDI. There were also no differences in alpha-diversity based on toxin presence measured by EIA or activity measured by Vero-cell rounding. Although toxin ELISA and Vero-cell positivity reflected the increased abundance of *C. difficile* in the sample, prior studies have demonstrated that asymptomatic colonized patients can have a high relative abundance of *C. difficile* present yet remain asymptomatic,^32^ again suggesting the role of non-*C. difficile* related factors that impact symptom status. Toxin presence continues to be used in a variety of recommended testing algorithms, and prior studies done in adults have demonstrated the ability of a positive toxin test to predict disease severity and recurrence.^31^ However, fulminant CDI has occurred in the setting of a negative toxin text^33^. Due to the similarities in toxin presence between asymptomatic and symptomatic pediatric patients with positive *C. difficile* tests, we elected to evaluate the intestinal microbiome as a potential contributor.

Unlike our study of *C. difficile* toxins, we identified significant differences in the intestinal microbiome and associated metabolites that effectively differentiated the two conditions. The results from our study suggest that the intestinal microbiome may play a critical role in symptom development. This is consistent with the observation that exposure to broad-spectrum antibiotics, with their prominent disruption of the intestinal microbiome, is the leading risk factor for CDI.

In our cohort, participants with asymptomatic colonization had a higher relative abundance of common gut commensals such as *Faecalibacterium* (q=0.041), *Blautia* (q=0.041), and *Bifidobacterium* (q=0.063) than those with symptomatic disease. Patients with symptomatic CDI had a higher relative abundance of Proteobacteria phyla and *Escherichia*/*Shigella* genera. Crobach et al. previously evaluated differences in adult patients with CDI, those with asymptomatic colonization, and healthy controls and reported that patients with CDI had higher relative abundance of *Bacteroides* and *Veillonella* and lower abundance of genera belonging to the Ruminococcaceae family and Actinobacteria phylum. Although this study did not report metabolites, the manuscript noted that many lower-abundance genera in those with CDI were known short-chain fatty acid (SCFA)-producers, including butyrate. Zhang et al. also compared gut microbiota between eight adult patients with CDI and eight adult patients with asymptomatic colonization.^9^ Although smaller numbers than our study, they confirmed similar findings with the gut microbiota of patients with CDI consisting of a greater proportion of Proteobacteria and a smaller proportion of Bacteriodetes and Firmicutes than those with asymptomatic *C. difficile* colonization. They also demonstrated a significantly higher proportion of *Escherichia*/*Shigella* in CDI patients vs asymptomatic carriers (23.9% versus 8.7%, P<0.05), as demonstrated in our pediatric cohort. Zhoe et al. performed a longitudinal study of adult ICU patients with *C. difficile* colonization and CDI and noted an increase in the relative abundance of SCFA and lactic acid-producing bacteria when patients transitioned from *C. difficil*e negative to *C. difficile* positive. The authors hypothesized that their findings suggest that the intestinal microbiota may react to *C. difficile* colonization by eliciting potentially protective responses such as an increased abundance of SCFA-producing bacteria.

Importantly, the trends in our gut microbiome dataset, with a higher abundance of members of the Proteobacteria phyla and a lower abundance of members of the Firmicutes phyla, were maintained even when the gut microbiota was analyzed within specific patient cohorts such as IBD or cancer. In fact, symptom status was the most significant driver of the intestinal microbiome even when considering various patient factors including medication use, comorbidity profile, age, and sex. Microbial associations with symptom status were also not sensitive to which patient factors were adjusted for in the analyses. These together suggest that the intestinal microbiome is strongly and consistently associated with *C. difficile* symptom status, even among patients with different comorbidities.

Similar to the Crobach et al. study, phyla, families, and genera classically associated with butyrate production had significantly higher relative abundance in children with asymptomatic colonization in our study. Concordantly, butyrate was higher in stool samples from children with asymptomatic colonization than those with symptomatic disease, suggesting a potential protective role. Short-chain fatty acids (SCFAs), including butyrate, are the major metabolic product of fiber metabolism by the intestinal microbiome and have demonstrated an increasingly important role in intestinal inflammation.^10^ Butyrate has demonstrated the ability to improve intestinal barrier function and mucosal immunity and has been the focus of increasing study in CDI. Butyric acid can decrease intestinal permeability and enhance colonic defense barriers by increasing mucin production and antimicrobial peptide levels, thus preventing the development of infection.^34^ Therefore, depletion of butyrate-producing bacteria could lead to epithelial dysfunction and a higher osmotic load in the intestinal lumen, leading to the diarrhea seen in CDI.^9^ The role of butyrate has been studied in a variety of animal and *in vitro* model systems. Fachi et al. demonstrated that butyrate supplementation could protect against CDI in mice, which was not dependent on alterations in *C. difficile* colonization or toxin production but rather a direct effect on intestinal epithelial cells.^13^ Specifically, butyrate increased the resistance of intestinal epithelial cells to *C. difficile* toxins by increasing and stabilizing hypoxia-inducible factor 1_α_ (HIF-1_α_). HIF-1_α_ reduces intestinal epithelial damage and improves the immune response; both relevant factors to attenuate the intestinal inflammation characteristic of symptomatic CDI. Butyrate has also been shown to inhibit the growth of diverse *C. difficile* strains *in vitro*.^28^

Of note, other studies have demonstrated alternative metabolic pathways that may be relevant in *C. difficile* colonization and CDI. Fishbein et al. demonstrated that asymptomatically colonized patients’ microbiomes were enriched in Clostridia species and had altered carbohydrate metabolism. However, the asymptomatic cohort in this study was defined differently than in our study and included adult patients with diarrhea who tested negative for toxin by EIA. As previously demonstrated, the presence or absence of toxins does not adequately differentiate colonization from disease in our pediatric cohort. The Fishbein et al. study may be evaluating a slightly different clinical scenario, i.e., comparing patients who are symptomatic but toxin-negative versus symptomatic and toxin-positive, which would explain the discrepancies in results between our studies.

Our study has some limitations. Patient samples from a single pediatric center may not represent the larger population, though we again note the consistency of identified microbiome-symptom associations across different disease conditions. Analysis of the presence and function of the gut microbiota was done via 16S rRNA sequencing and measurement of a single stool metabolite, butyrate. Therefore, the intestinal microbiome’s function, which can be more fully captured through metagenomics and metabolomics, cannot be fully detailed and should be addressed through future studies. Longitudinal studies with repetitive sampling would also allow for a more robust analysis of contributing factors.

In conclusion, through this study comparing the intestinal microbiome and butyrate of children with asymptomatic colonization versus CDI, we identify that the microbiome, and not toxin, can better differentiate between these two conditions. These data suggest that the presence of butyrate, as mediated by the intestinal microbiome, is a critical protective factor in the development of symptomatic CDI. Collectively, these data support that butyrate supplementation, or dietary manipulation that results in increased levels of butyrate-producing organisms, warrants further evaluation for their therapeutic potential for both prevention and treatment of CDI.

## Abbreviations

CDI: *Clostridioides difficile* infection
rRNA: ribosomal Ribonucleic Acid
ASV: Amplicon Sequence Variant

## Author’s contributions

MRN, BAS, MHS, EPS, and SRD contributed to the study design. MRN contributed to the sample collection. BAS contributed to the sample processing and sequencing, data processing, and statistical analysis. SM performed the final data processing, statistical analysis, and figures. MC, LZ, and MJM contributed to sample processing and analysis. ERG and MJM contributed to LC-MS/MS sample preparation, data collection, and analysis. MRN and SRD obtained the research funding supporting this study. MRN, SM, BAS, MHS, and SRD wrote the initial version of the manuscript, and all authors reviewed and approved the final version.

## Data Transparency Statement

Deidentified data will be shared upon request.

