## Supplemental Figure 1 for "The Gut Microbiome and Butyrate Differentiate *Clostridioides difficile* Colonization and Infection in Children"

**A**

| | ELISA + (N=28) | ELISA - (N=68) | P-value | VERO + (N=40) | VERO - (N=54) | P-value |
|---|---|---|---|---|---|---|
| **Symptom** | | | | | | |
| Asymptomatic | 11 (39.3%) | 36 (52.9%) | 0.32 | 20 (50.0%) | 25 (46.3%) | 0.88 |
| Symptomatic | 17 (60.7%) | 32 (47.1%) | | 20 (50.0%) | 29 (53.7%) | |
| **Age** | 6.5 (3-13) | 10 (5-15) | 0.088 | 9 (4-14) | 10.5 (5.25-16) | 0.18 |
| **Gender** | | | | | | |
| Male | 18 (64.3%) | 33 (48.5%) | 0.24 | 24 (60.0%) | 26 (48.1%) | 0.35 |
| Female | 10 (35.7%) | 35 (51.5%) | | 16 (40.0%) | 28 (51.9%) | |

**B** **C**

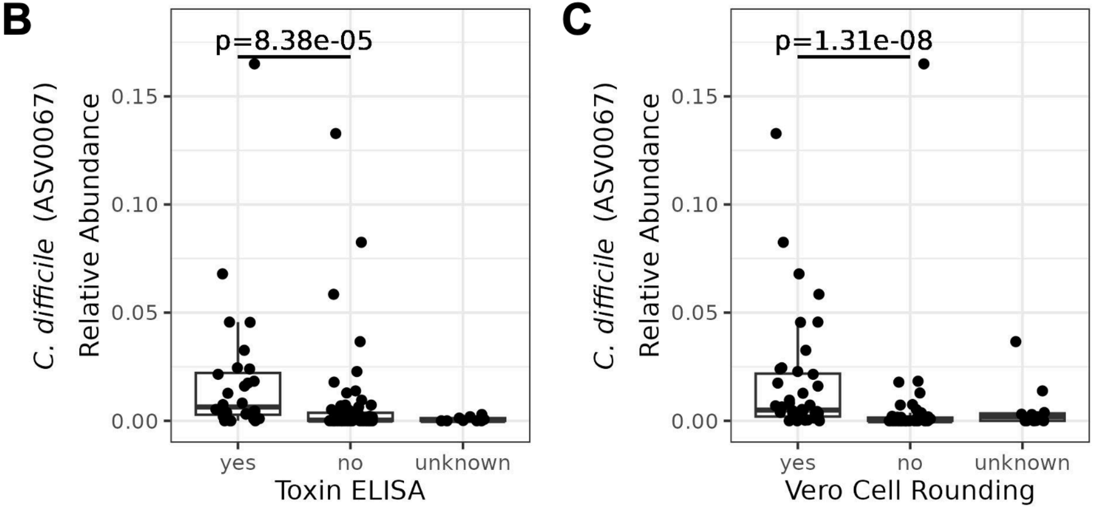


**D**

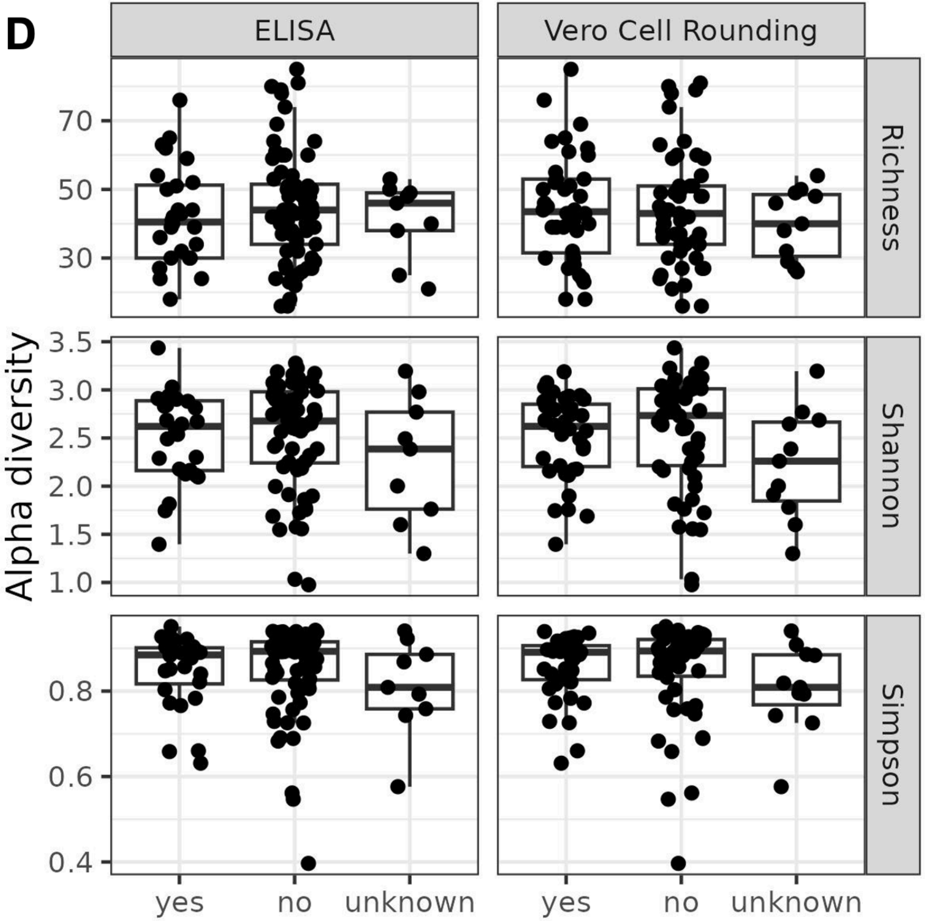
