## Supplementary figures and images for "The Gut Microbiome and Butyrate Differentiate *Clostridioides difficile* Colonization and Infection in Children"

### Supplemental Figure 2

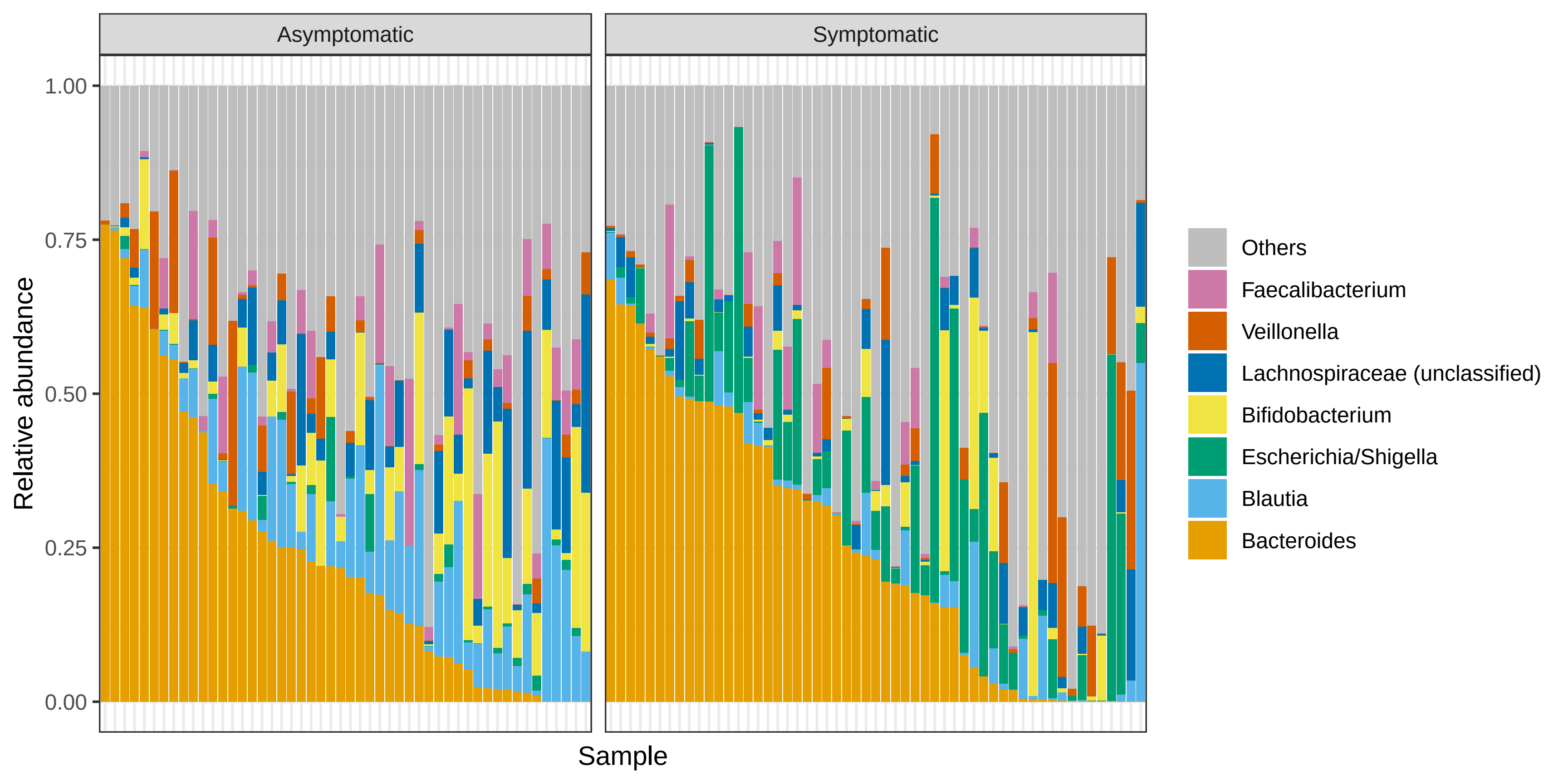

### Supplemental Figure 3

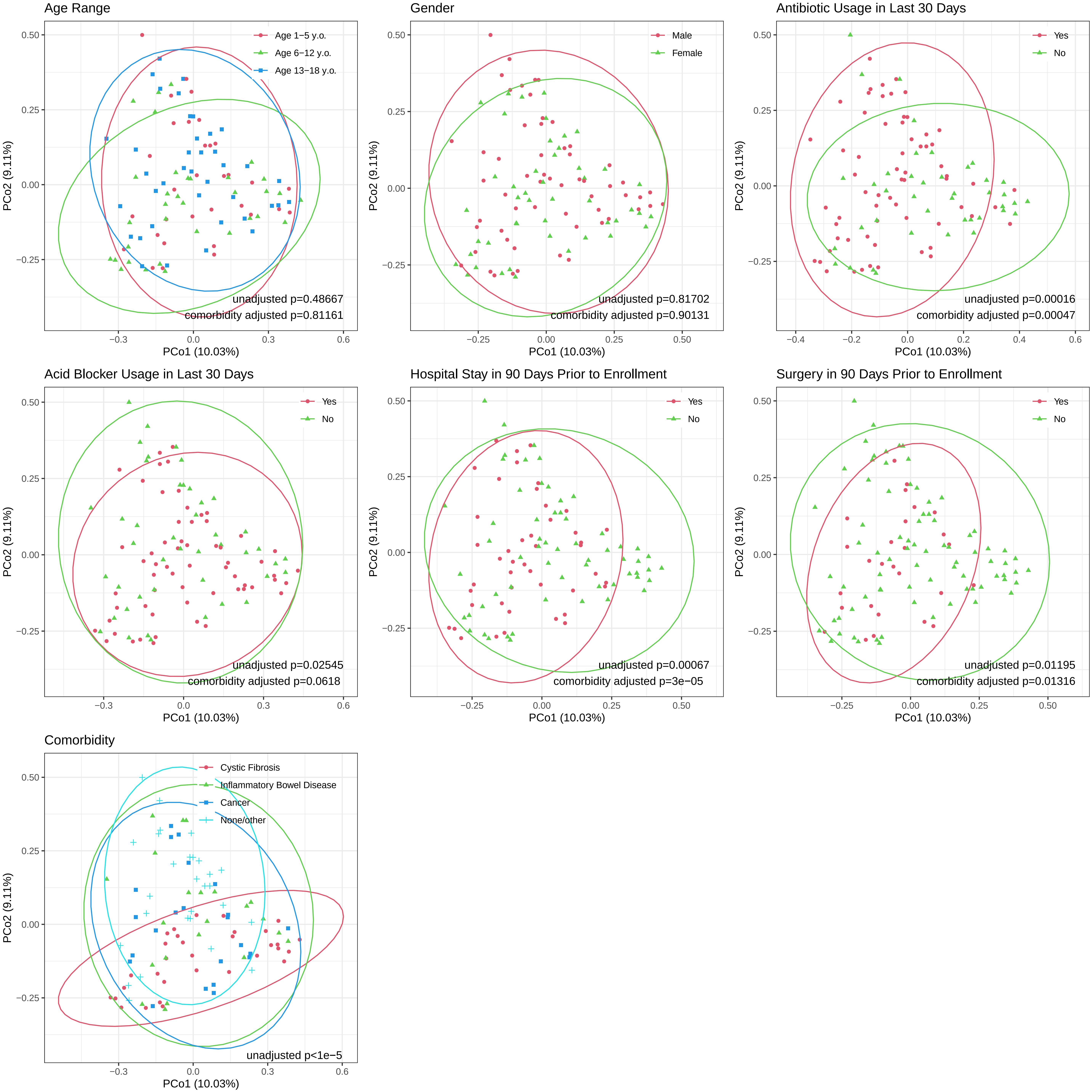

### Supplemental Figure 4

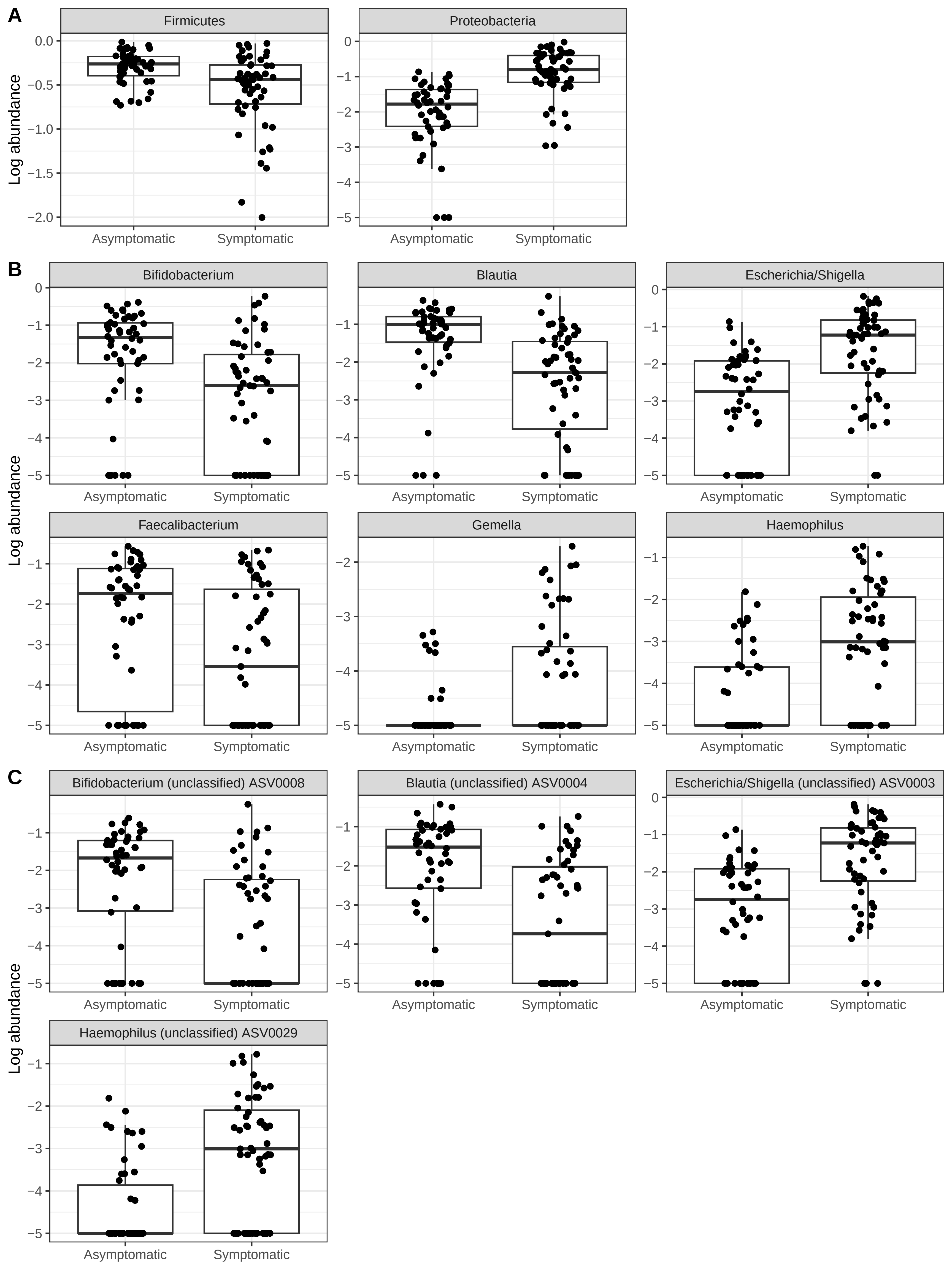

### Supplemental Figure 5

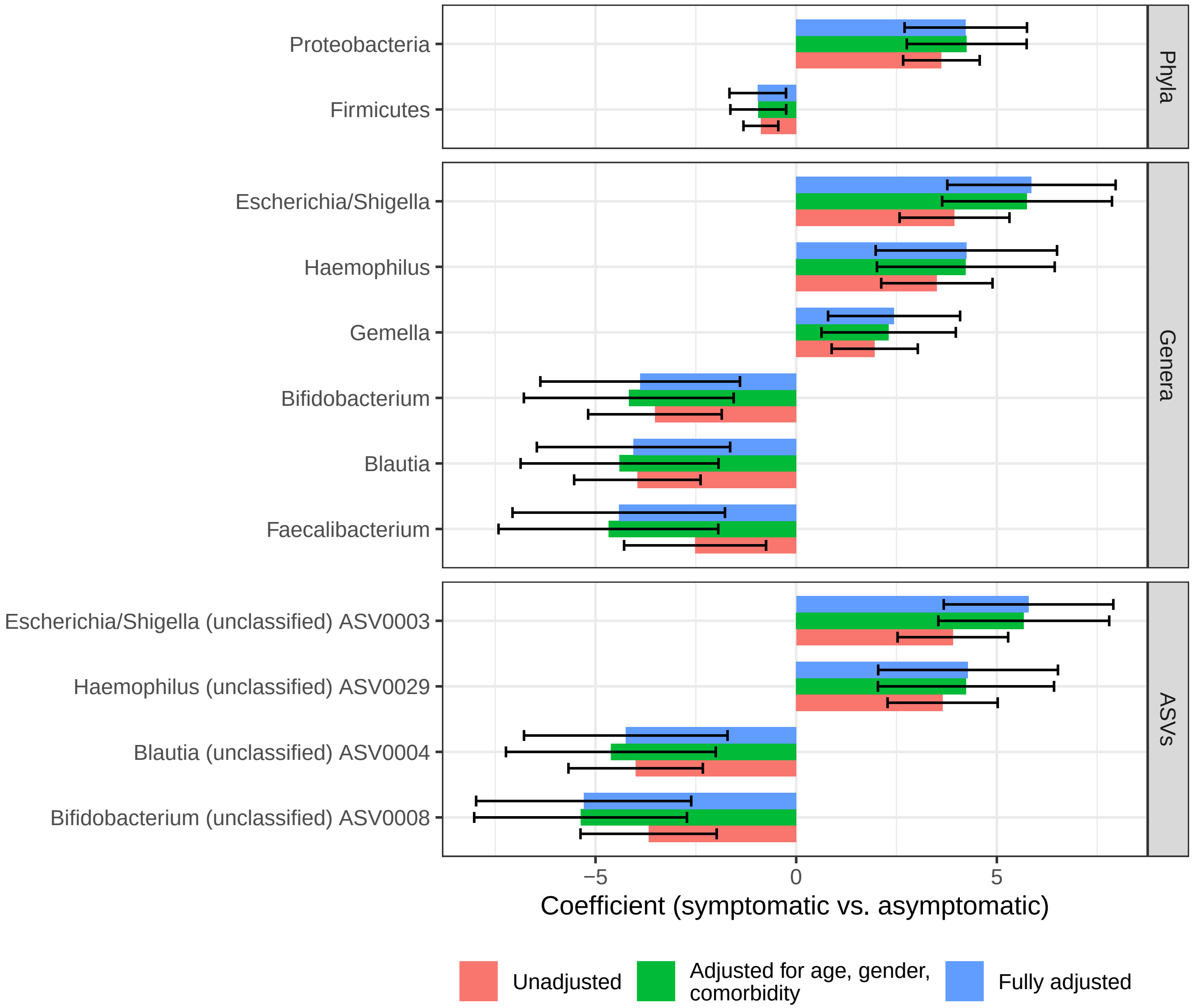

### Supplemental Figure 6

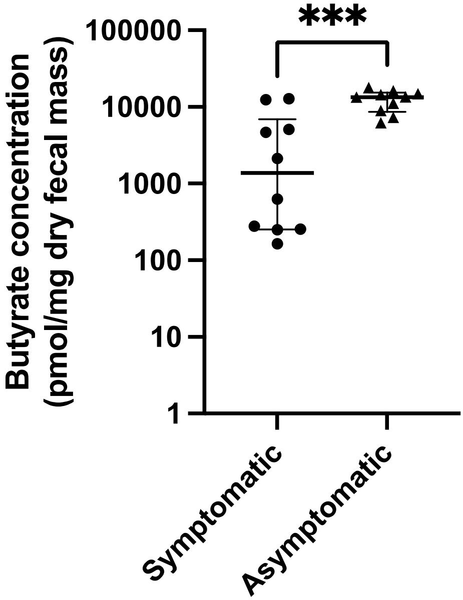
